# A broad cathepsin inhibitor blocks crystal-stimulated inflammasome-dependent and -independent inflammation, and gout arthritis

**DOI:** 10.1101/2024.07.01.601464

**Authors:** Laura Alejandra Ariza Orellano, Chunhui Zeng, Jiyun Zhu, Matthew Bogyo, Kenneth L. Rock, Jiann-Jyh Lai

## Abstract

In the disease gout, monosodium urate (MSU) crystals nucleate in joints and cause acute painful arthritis that can damage the affected joints. Similarly, the deposition of other crystals or irritant particles in tissues elicits an inflammatory response that can cause disease. These various particles stimulate macrophages to produce the proinflammatory cytokine interleukin 1β (IL-1β), which is a major driver of the ensuing inflammation. Here we show that in vivo and in vitro, broad spectrum cathepsin inhibitors, like VBY-825, blocked the activation of inflammasomes, which are known to be essential in generating bioactive IL-1β in response to crystals. In addition, the cathepsin inhibitors blocked an inflammasome-independent pathway that also generates mature IL-1β and which contributed substantially to crystal-stimulated inflammation in vivo. Through these effects, the cathepsin inhibitors markedly reduced gout arthritis and inflammation to the unrelated crystal silica, which is the etiologic agent in the disease silicosis. The cathepsin inhibitors didn’t affect any of the inflammatory processes after bioactive IL-1β was present in tissues. They also didn’t inhibit LPS-stimulated inflammation *in mice*, or TNF-⍺ production from macrophages. These findings provide proof of concept that cathepsin inhibitors are a novel class of anti-inflammatories that can inhibit crystal-stimulated disease with unique mechanisms of action.

## Introduction

When irritant crystals and particles deposit in tissues, they elicit an inflammatory reaction (1) A classic example of this process is the disease of gout (2). Gout occurs in hyperuricemic patients when uric acid periodically nucleates into monosodium urate (MSU) crystals. When this occurs in a joint, it causes severe arthritis that is exquisitely painful. The affected site becomes hyperemic, edematous, and infiltrated with neutrophils and monocytes/macrophages. Repeated bouts of this intraarticular inflammation can damage and disfigure the affected joint. MSU crystals can deposit in other sites, such as the kidney, and similarly cause tissue damage. Tissue deposition of many other crystals or particles similarly elicits inflammation and causes disease, e.g., silica (silicosis) (3), asbestos (asbestosis) (4), cholesterol (atherosclerosis) (5), and titanium particles (post-joint implant inflammation) (6).

One of the key mediators driving these particle-stimulated inflammatory responses is the proinflammatory cytokine interleukin-1β (IL-1β). Mice that genetically lack the receptor for IL-1 (IL-1R) or its signaling components, or the IL-1 cytokines have a markedly attenuated inflammatory response to particles, including MSU crystals (7-9), also Supplemental Figure 1. Similarly, when human patients experience an acute gouty flare, treating them with an anti-IL-1β antibody (canakinumab) or an IL-1 competitive antagonist (IL-1Ra) rapidly ameliorates their inflammatory symptoms (8, 10, 11).

IL-1β is initially synthesized as a longer polypeptide pro-form, which is not biologically active (12). Pro-IL-1β needs to be cleaved to generate the mature and biologically active form of IL-1β. The protease that makes this activating cleavage in cells is caspase 1 (13, 14). Caspase 1 is synthesized as a zymogen. Its autoactivation is controlled by pattern-recognition receptors, such as certain NOD-like receptors. In the case of crystals and particles, the key NOD-like receptor is NLRP3 (15, 16). Upon exposure to activating stimuli, such as crystals, NLRP3 in cells is stimulated to associate with the scaffolding protein ASC, the latter of which associates with procaspase 1 and polymerizes into a macroscopic filamentous structure (aka specks) (17). This process leads to the autocleavage of caspase 1 into its catalytically active form. The NLRP3-poly-ASC-caspase 1 complex is called the NLRP3 inflammasome.

There are multiple different mechanisms that can lead to the stimulation of the NLRP3 inflammasome response (18, 19). In the case of crystals, this process starts when these particles are ingested by macrophages through phagocytosis. Inhibiting this internalization of crystals blocks the activation of the NLRP3 inflammasome (20). Some of the crystal-containing phagosomes subsequently rupture and such rupture, even without crystals, is sufficient to activate the NLRP3 inflammasome (20). In this situation, it is thought that one of the things that NLRP3 senses is the release of proteases into the cytosol. It was initially thought that the key protease was cathepsin B (21), but subsequent studies showed that many different cathepsins could initiate this process, i.e., there was redundancy (22).

In cell culture, caspase 1 and the inflammasome are absolutely required for macrophages to make active IL-1β in response to crystals (23). However, in vivo, there is a substantial inflammasome-independent pathway of IL-1β production (24, 25). In this later situation, it is thought that proteases released from activated leukocytes, such as neutrophils, cleave extracellular pro-IL-1β that was released into the extracellular space (26). These leukocyte proteases are initially produced as zymogens, which can be cleaved into an active form by cathepsins (26).

Given the role of cathepsins in both crystal-stimulated inflammasome-dependent and inflammasome-independent inflammation, these proteases have the potential to be unique therapeutic targets to block such responses. Here we test this hypothesis with broadly active small molecule cathepsin inhibitors.

## Results

### A broad spectrum cathepsin inhibitor blocks crystal-stimulated IL-1β production in cultured cells

VBY-825 is a small molecule ketoamide that is a potent reversible inhibitor of multiple cysteine cathepsins, including cathepsins B, L, S, V, F, and K (27). We found this agent also inhibits cathepsin C, but with lower potency (Supplemental Figure 2). VBY-825 inhibits cathepsins by forming a reversible covalent hemithioacetal bond with their catalytic sites’ cysteines. VBY-825 is able to inhibit cathepsins in living cells in vitro (27)

To investigate the effect of VBY-825 on IL-1β responses to crystals, we stimulated LPS-primed naïve (resident) peritoneal macrophages in vitro with silica or MSU crystals in the presence or absence of the inhibitor. VBY-825 reduced the IL-1β responses to both crystals (Figure 1, A and B). The magnitude of this inhibition was greater for silica than for MSU, but both reductions were significant. Similar results were obtained when using another broad spectrum cathepsin inhibitor, CA-074 methyl ester (CA-074 Me) (Figure 1A). At the concentrations used in these experiments, CA-074 Me also broadly inhibits multiple cysteine cathepsins (22). MSU and silica crystals cause macrophages to die via pyroptosis (28, 29), which can be measured by the release of LDH, and VBY-825 inhibited this process (Figure 1C). In contrast, the LPS-stimulated production of TNF-α from the same experimental groups, which is known to be NLRP3-independent, was unaffected by VBY-825 (Figure 1D and Supplemental Figure 3). These latter results indicated that VBY-825 was not generally inhibiting the ability of MØs to produce cytokines.

**Figure 1.**
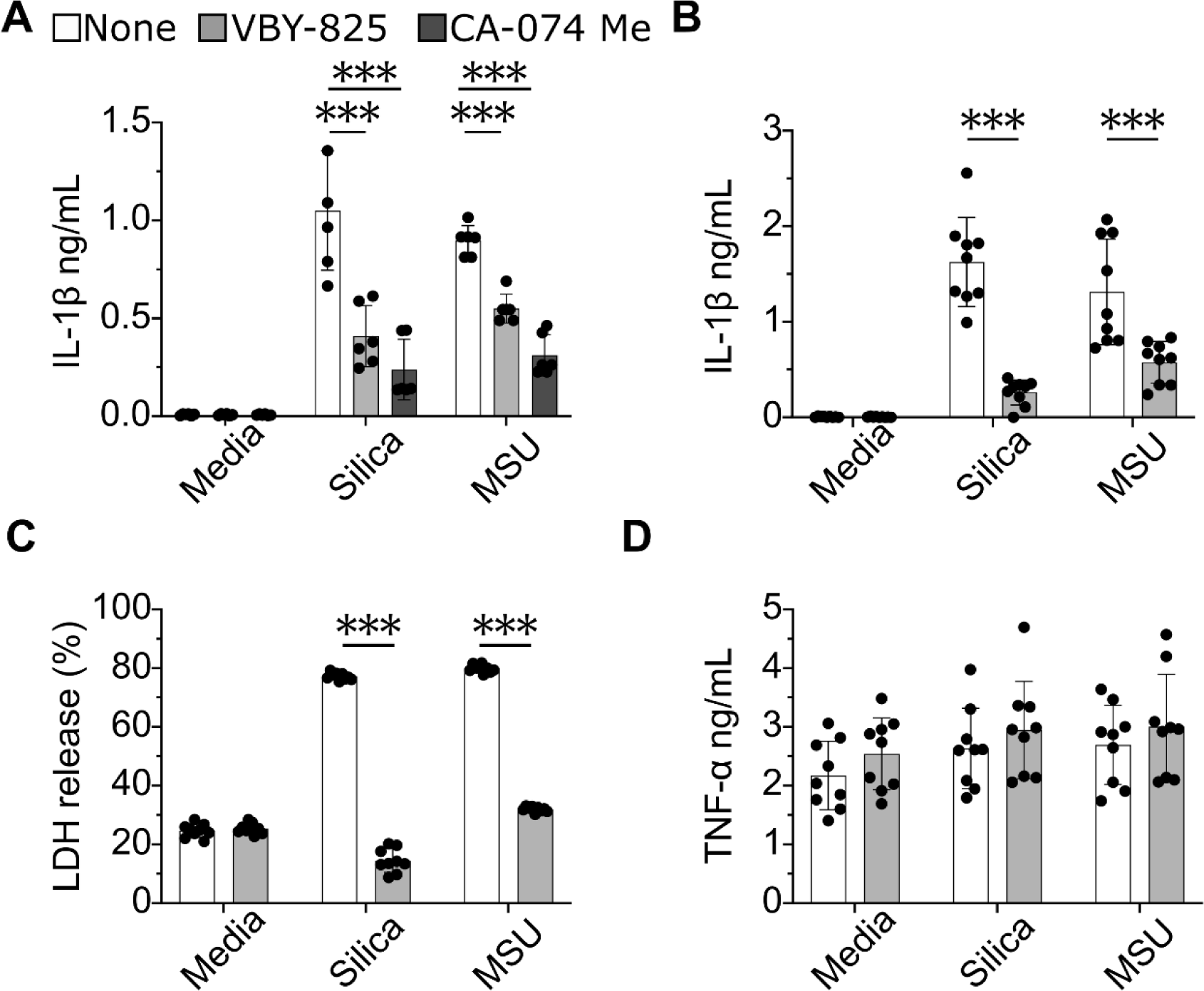
Broad spectrum cathepsin inhibitors block crystal-stimulated IL-1β production in macrophages. (**A**) Murine resident peritoneal macrophages (RPMs), 1x10^5^/well, were primed with LPS (200 ng/mL) for 2 hours, followed by vehicle, VBY-825 (120 µM) or CA-074 Me (5 µM) for another hour, and then media, Silica (15 µg/well), or MSU (25 µg/well) was added and cells were incubated for another 4 hours. IL-1β in the culture supernatant was measured by ELISA. (**B-D**) RPMs, 1x10^5^/well, were incubated with LPS (200 ng/mL) for 2 hours, followed by another 2 hours with or without VBY-825 (120 µM), and then stimulated for 4 hours with or without Silica (15 µg/well) or MSU (25 µg/well). Supernatants were collected to measure (**B**) IL-1β, (**C**) LDH, and (**D**) TNF-α. Data are combined from three independent experiments with the same parameters and shown as mean ± SD. ***p<0.001.

### VBY-825 inhibits crystal (MSU) stimulated inflammation in vivo

Previous studies have shown that administering a single dose of VBY-825 in vivo inhibits cathepsins for over 24 hours (27). Therefore, we next investigated whether VBY-825 could inhibit crystal-induced inflammation in vivo. Mice were injected with VBY-825 i.v. and then challenged with crystals i.p. After 4 hours, the peritoneum was lavaged, and the numbers of neutrophils and macrophages were quantified by immunofluorescent staining and flow cytometry. The injection of MSU crystals stimulated an influx of neutrophils and macrophages into the peritoneum, as expected (Figure 2, A and B). In contrast, in animals treated with VBY-825, the MSU-induced peritonitis was markedly attenuated (Figure 2, A and B). The magnitude of the reduction in MSU-stimulated inflammation from VBY-825 treatment was similar to that caused by genetic loss of the IL-1R (Supplemental Figure 1). To test whether these findings would extend to another unrelated crystal, we also injected mice with silica and found that VBY-825 similarly inhibited the ensuing inflammation (Figure 2, C and D). Therefore, VBY-825 inhibited inflammation of multiple unrelated and structurally distinct crystals in vivo. Similar results were obtained treating the animals with CA-074 Me (Figure 2, E and F) indicating that the anti-inflammatory effects of these agents were almost certainly due to cathepsin inhibition.

**Figure 2.**
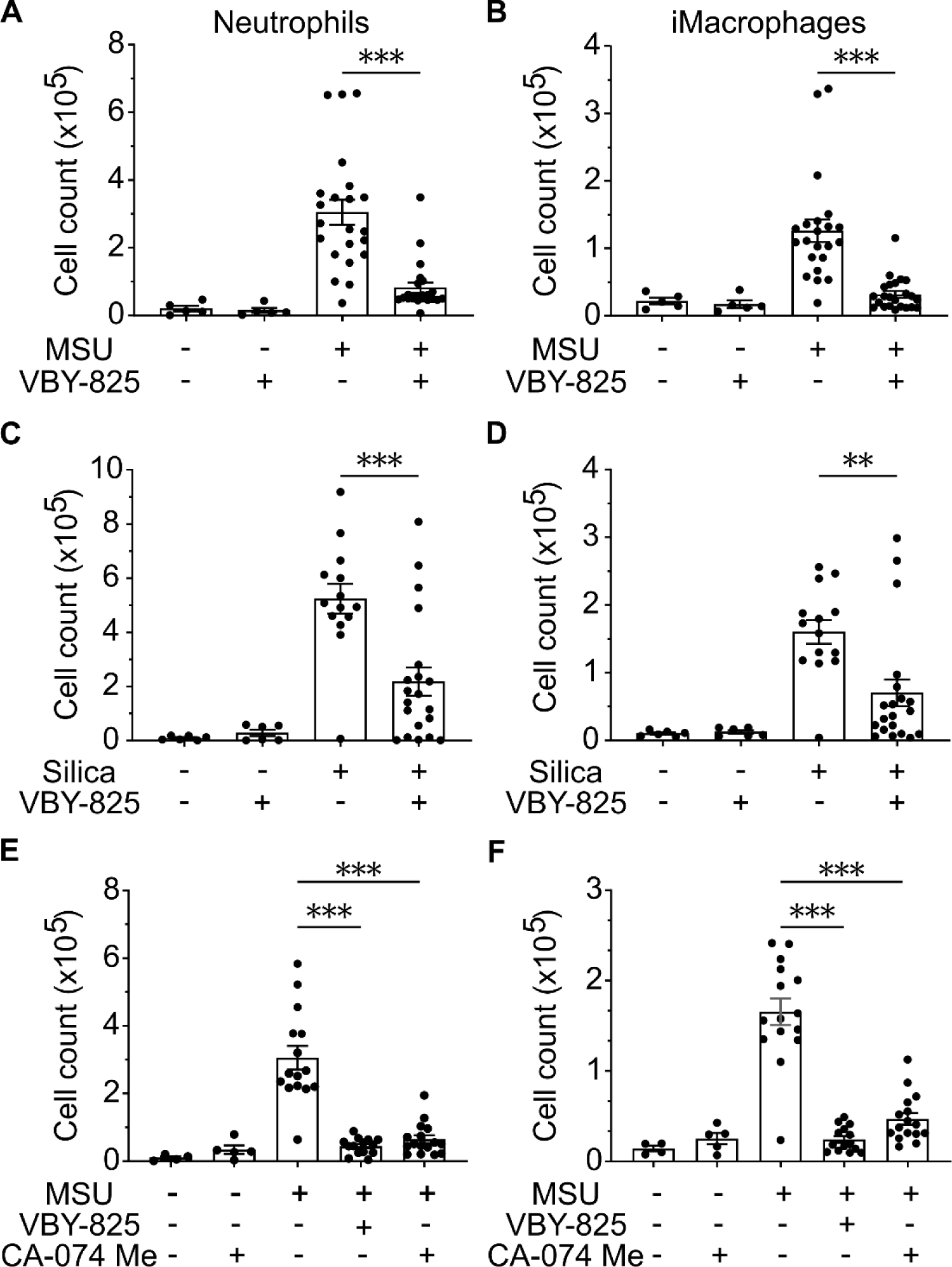
Cathepsin inhibitors reduce crystal-stimulated inflammation *in vivo*. Female B6 mice were i.v. injected with or without VBY-825 (10 mg/Kg) 1 hour prior to i.p. injection with (**A and B**) MSU (0.1 mg/mouse), (**C and D**) Silica (0.1 mg/mouse), or PBS as the negative control. Four hours later, the numbers of infiltrating neutrophils and inflammatory macrophages (iMacrophages) in the peritoneum were counted by flow cytometry. (**E and F**) Similar to (A), except mice were treated with or without VBY-825 (10 mg/Kg) or CA-074 Me (10 mg/Kg). Data are combined from 3 independent experiments with the same parameters and represented as means ± SEM. **p<0.01; ***p<0.001.

### Mechanisms of VBY-825 anti-inflammatory effects

We next sought to determine how VBY-825 inhibited inflammatory responses in vivo. Given the above findings, our leading hypothesis was that VBY-825 was limiting the production of bioactive IL-1β. This hypothesis made the testable prediction that VBY-825 should affect a step involved in the production of bioactive IL-1β but not in events thereafter. To test this prediction, we evaluated the effect of VBY-825 on inflammation induced by injection of mature IL-1β. IL-1β administered directly into the peritoneum induced acute inflammation, with the recruitment of neutrophils and macrophages (Figure 3, A and B). Remarkably, co-administration of VBY-825 did not inhibit this response, while in the same experiment, it inhibited crystal-induced peritonitis (Figure 3, A and B). A similar result was obtained with CA-074 Me (Figure 3, C and D). These are important results because they demonstrated that the cathepsin inhibitors were not inhibiting steps in the inflammatory response that are downstream of IL-1β, e.g., neutrophil trafficking and diapedesis. These results pointed to the cathepsin inhibitors likely acting on a mechanism involved in the generation of bioactive IL-1β.

**Figure 3.**
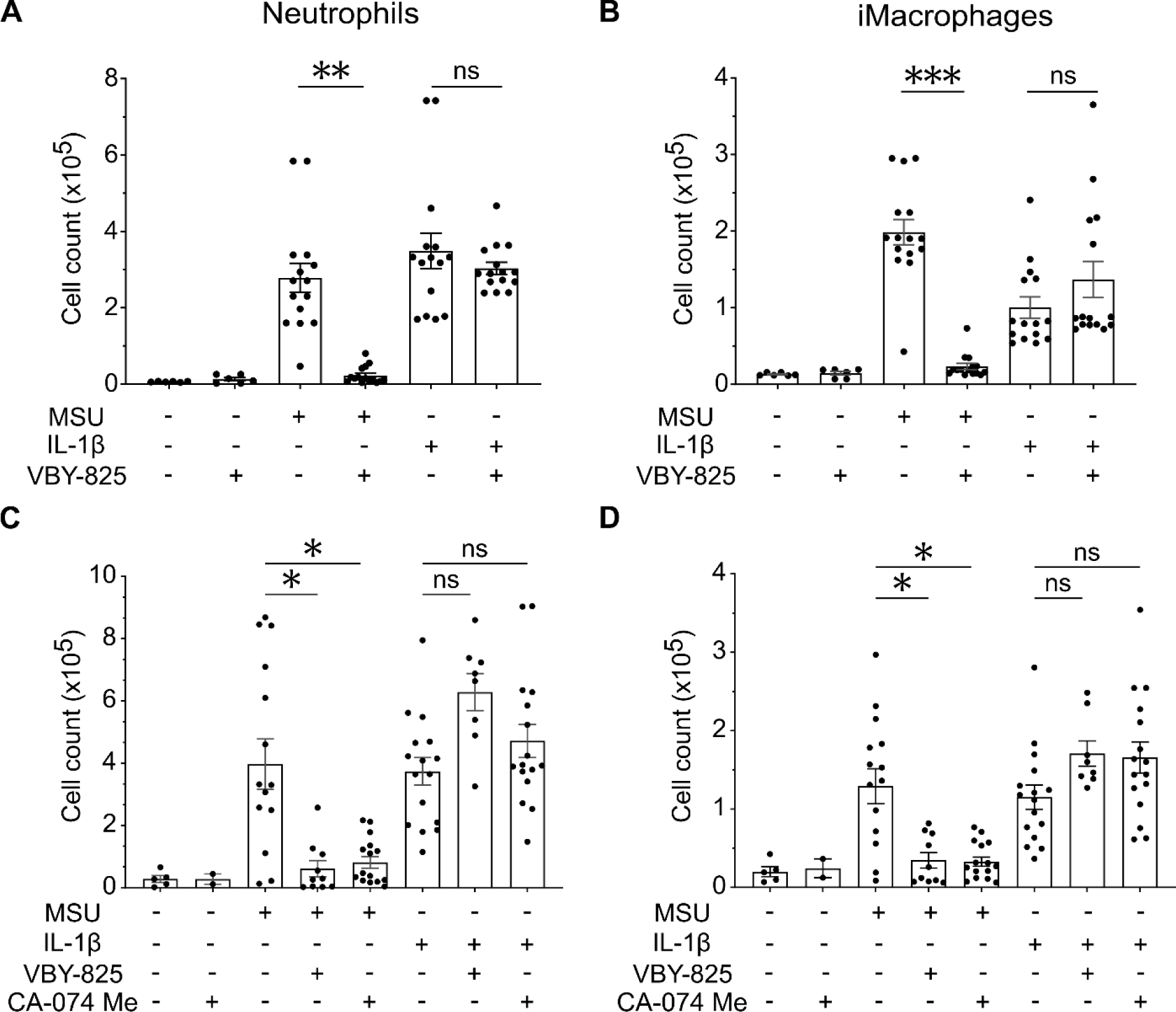
Cathepsin inhibitors block the pathway upstream of IL-1β. Female B6 mice were pre-treated with VBY-825 (i.v. 7.5mg/Kg) 1 hour prior to i.p. injection with or without MSU (0.1 mg/mouse) or IL-1β (1 ng/mouse) for another 4 hours, followed by peritoneal lavage to stain and count the infiltrating (**A**) neutrophils and (**B**) iMacrophages by flow cytometry. (**C and D**) Similar to (A and B), except mice were i.v. pre-treated with VBY-825 (7.5 mg/Kg) or CA-074 Me (10 mg/Kg) 1 hour prior to i.p. injection with or without MSU (0.1 mg/mouse) or IL-1β (1 ng/mouse) for another 4 hours. The data were combined from 3 independent experiments and represented as means ± SEM. *p<0.05; **p<0.01; ***p<0.001.

Cathepsins have been implicated in the activation of NLRP3 inflammasomes, which are required for the intracellular generation of mature IL-1β (20). To investigate whether VBY-825 was inhibiting this activation, we examined whether it blocked the formation of polymerized inflammasomes. To assay this event, we took advantage of a mouse model that expressed an ASC-fluorescent protein (citrine) fusion (30). When primed macrophages from these mice were stimulated with crystals in vitro, their inflammasomes polymerized into fluorescent specks, which could be quantified by imaging flow cytometry (Figure 4). Treatment with VBY-825 inhibited the formation of these specks in response to stimulation with crystals (Figure 4, A and B). When MSU crystals were injected i.p. into the ASC-citrine mice, macrophages containing fluorescent specks were present in the peritoneum. However, this production of specks in response to MSU crystals was inhibited when mice were treated with VBY-825 (Figure 4, C and D). These results demonstrated that VBY-825 was inhibiting the activation of NLRP3 inflammasomes *in vivo*.

**Figure 4.**
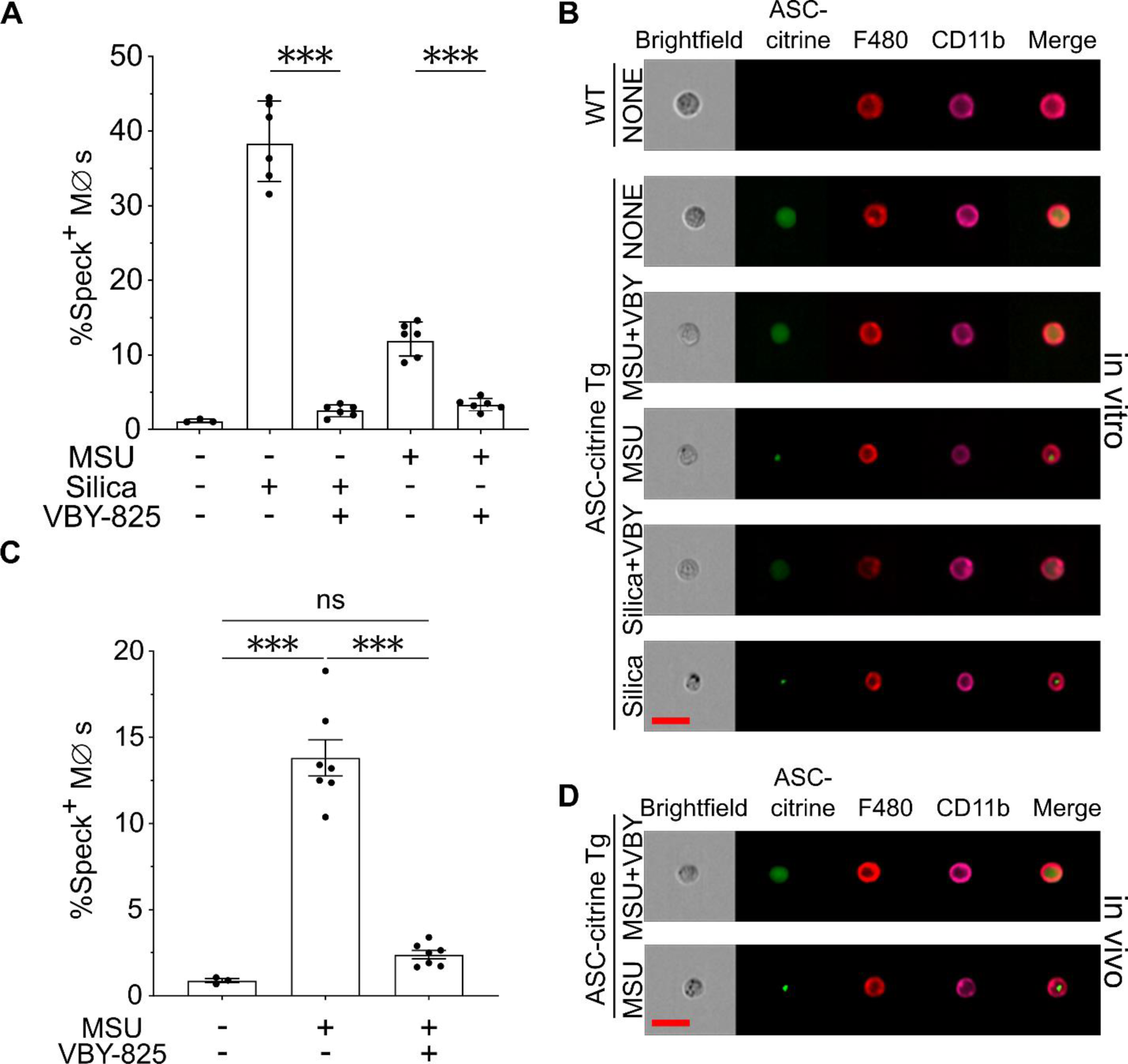
VBY-825 inhibits crystal-stimulated inflammasome activation *in vivo*. RPMs (10^6^ cells) from WT or ASC-citrine transgenic mice were primed with LPS for 3 hours, followed by treatment with 120 µM VBY-825 or vehicle for 1 hour. Afterward, the macrophages were stimulated with or without MSU (100 µg) or Silica (60 µg) for another 3 hours before staining them with anti-CD11b, anti-F4/80, and DAPI. The cells were then analyzed by imaging flow cytometry, and (**A**) Percentage of speck^+^ macrophages, and (**B**) Representative FlowSight images are presented. (**C and D**) Female ASC-citrine mice were pre-treated i.v. with VBY-825 (7.5mg/Kg) or vehicle for 1 hour, followed by i.p. injection with or without MSU (0.1 mg/mouse), and 4 hours later peritoneal exudative cells (PECs) were collected, stained, and analyzed by imaging flow cytometry. (**C**) The percentage of speck^+^ cells was calculated within resident macrophages (CD11b^hi^ F480^hi^) population, and (**D**) Representative FlowSight images are presented. The data were combined from 3 experiments and represented as means ± SD (A) or mean ± SEM (C). ***p<0.001. Red bar=20 µm.

We also examined the effect of VBY-825 on the inflammasome-independent pathway of inflammation. For these experiments, we used caspase 1-null mice. Without caspase 1, cells lack functional NLRP3 inflammasomes, and therefore any inflammation that occurs is independent of this mechanism. These mutant mice were injected i.p. with MSU crystals. Despite the absence of functional NLRP3 inflammasomes, caspase 1/11 deficient mice still generated an inflammatory response to MSU crystals (31). Importantly, VBY-825 also inhibited this inflammasome-independent response (Figure 5). Taken together, these results demonstrate the VBY-825 inhibited both NLRP3 inflammasome-dependent and -independent crystal-induced inflammatory responses in vivo.

**Figure 5.**
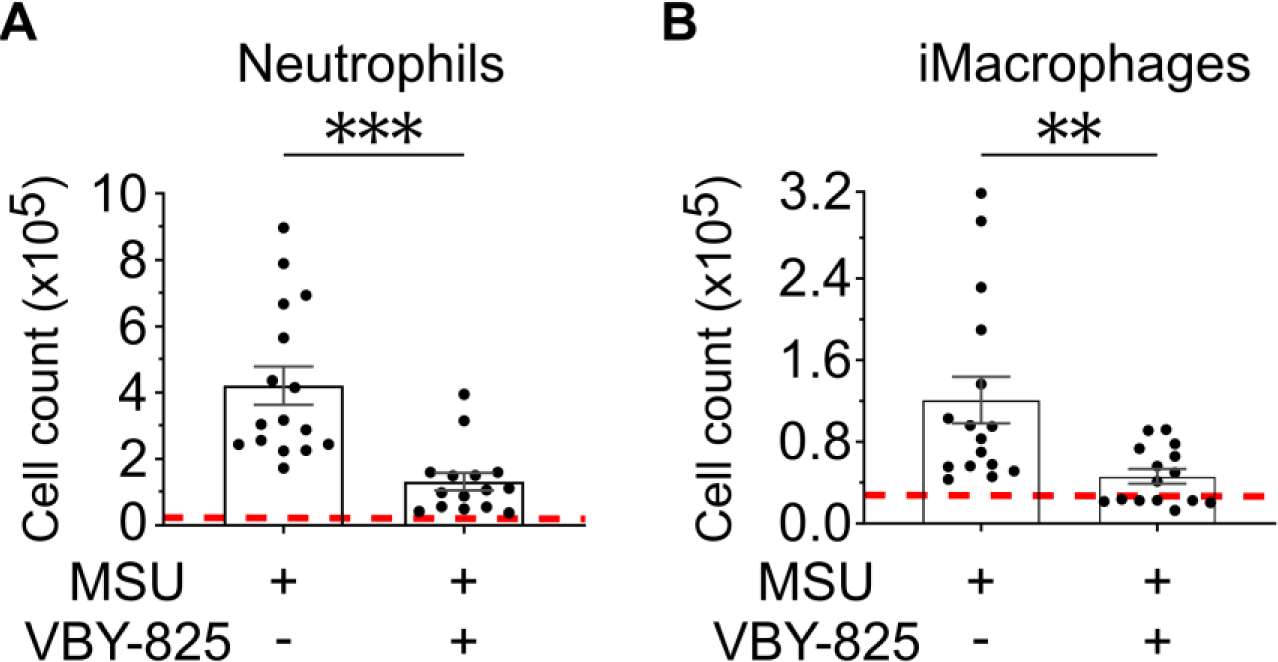
VBY-825 inhibits the crystal-stimulated inflammasome-independent component of inflammation *in vivo*. Caspase1/11 double-knockout (Casp1/11 DKO) mice were pre-treated i.v. with or without VBY-825 (7.5 mg/Kg) for 1 hour, followed by i.p. injection with or without MSU (0.1 mg/mouse), and 4 hours later (**A**) neutrophils and (**B**) inflammatory macrophages in the peritoneal lavage were analyzed as described in Figure 2. Red dashed line = background (no MSU stimulation). The data were combined from 3 experiments and represented as means ± SEM. **p<0.01; ***p<0.001.

### VBY-825 reduces inflammation in gout arthritis

In a final set of experiments, we investigated whether VBY-825 could reduce inflammation in a preclinical mouse model of gout. In this system, MSU crystals versus saline were injected into the intraarticular space of the knee. This model resembles the spontaneous crystal deposition that occurs in the intraarticular space of joints in human gout. The ensuing inflammation was then quantified by injecting luminol and measuring bioluminescence in the knee using an IVIS imaging system. Luminol is a redox-sensitive small molecule that bioluminesces in response to reactive oxygen species generated by myeloperoxidase in neutrophils and monocytes (32, 33). Bioluminescence was significantly elevated in the joints that received MSU crystals, indicating the presence of activated inflammatory cells. Remarkably, treatment with VBY-825 i.p. significantly reduced the MSU-stimulated bioluminescence signal (Figure 6, A and B).

**Figure 6.**
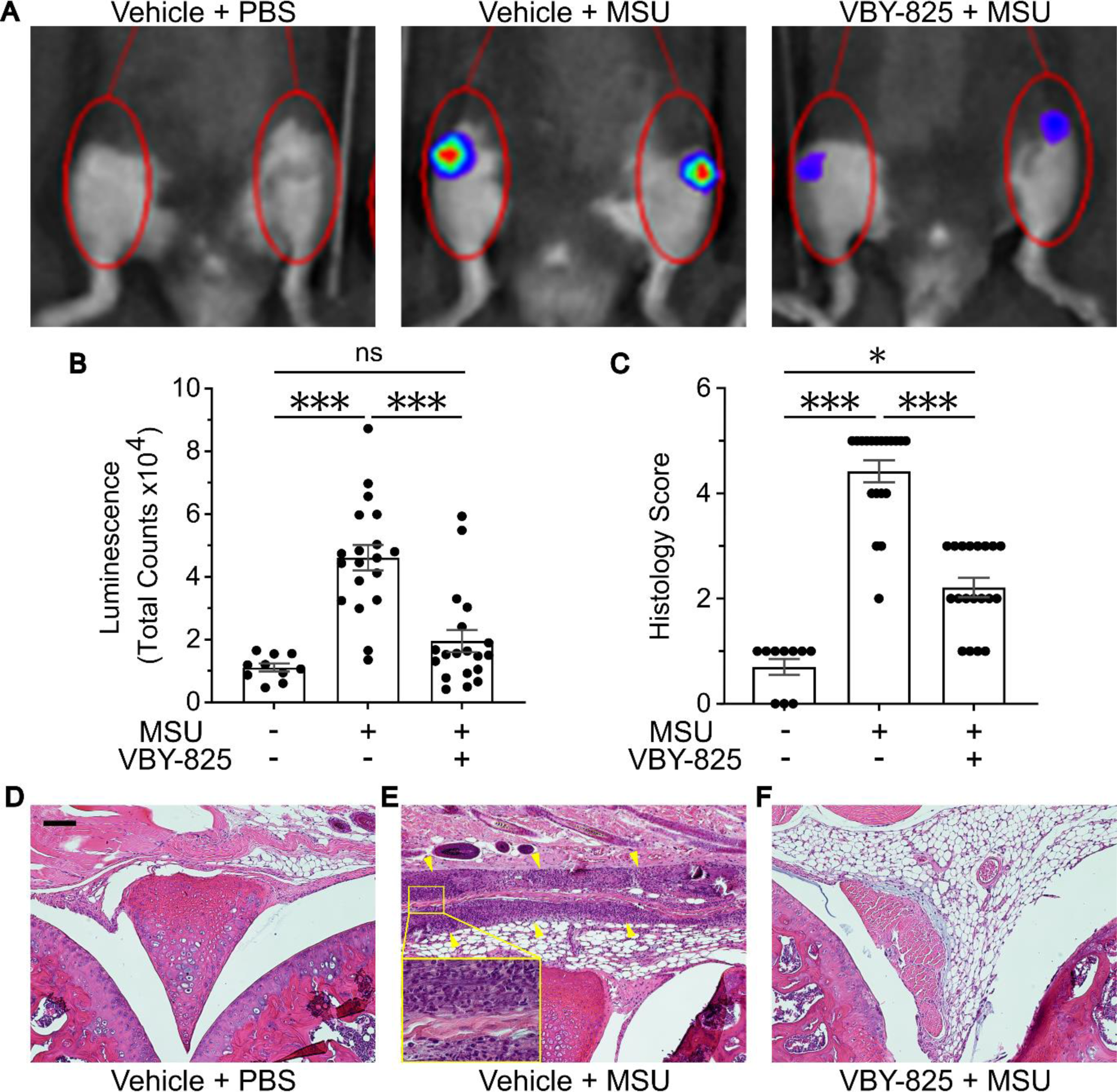
VBY-825 reduces inflammation in MSU-induced arthritis. Male B6 mice were i.p. injected with VBY-825 (1 mg/mouse) or vehicle 1 hour prior to intra-articular injection of MSU (150 µg/10µL) or PBS. Twenty-four hours after MSU stimulation, Luminol (200 mg/Kg) was i.v. injected, followed by bioluminescent imaging using an IVIS instrument. (**A**) Representative IVIS images of bioluminescence in the various treatment groups. (**B**) Intensity of bioluminescence in the knees of animals injected with or without MSU ± VBY-825. (**C**) The histology score of inflammation in the H&E sections was assessed by 3 people and averaged for each knee. (**D-F**) Representative pictures of knee sections (H&E stain) from the various groups. Scale bar=100 μm. The area between the top and bottom yellow arrows in (E) contains intense neutrophil infiltration. Insert in (E) is the highlighted region at higher magnification. The data were combined from 3 experiments and represented as means ± SEM. *p<0.05, ***p<0.001.

The above findings indicated that VBY-825 was reducing the amount of myeloperoxidase in joints containing deposits of MSU crystals. It seemed likely that this was due to a reduction in the inflammatory infiltrates in these sites. To determine if this was the case, we performed a histopathological examination of the injected joints and used an established system to score the inflammation (34). In knees injected with PBS, there were no inflammatory infiltrates, as expected (Figure 6D). In contrast, in the knees that received MSU crystals, there were infiltrates of neutrophils and mononuclear leukocytes (Figure 6E). However, in mice that also received VBY-825, these infiltrates were significantly reduced (Figure 6F). Together these findings indicated that VBY-825 suppressed the acute inflammation in gout arthritis.

## Discussion

Our major findings are that cathepsin inhibitors are a novel class of anti-inflammatory agents that can effectively treat gout arthritis and other crystal-stimulated inflammation with mechanisms of action that are different from currently available therapeutics. MSU and other crystals stimulate IL-1β-dependent inflammation, which is the underlying driver of the symptoms and pathology in gout arthritis and other crystal diseases such as silicosis (35). Two distinct pathways contribute to the crystal-stimulated production of bioactive IL-1β in vivo: inflammasome-dependent and inflammasome-independent (36, 37). We found that small molecules that inhibited multiple cathepsins, such as the ketoamide VBY-825, blocked both of these pathways and thereby effectively suppressed the crystal-stimulated acute neutrophilic inflammation in vivo in joints and elsewhere. Consistent with these results, VBY-825 reduced the level of MSU-induced inflammation down to that observed in a mouse that can’t respond to IL-1 (IL-1R KO). The cathepsin inhibitors didn’t affect inflammatory steps downstream of IL-1β production nor inhibit bacterial PAMP-stimulated inflammation in vivo or the proinflammatory cytokine TNF-α in vitro, and as such are selective rather than broad suppressors of inflammation.

MSU and Silica crystals stimulate NLRP3 inflammasome-dependent processing of pro-IL-1β into mature and bioactive IL-1β in macrophages. Cathepsins inhibitors have been shown to block this process in vitro (20, 22, 28, 38). Here we showed that the VBY-825 cathepsin inhibitor similarly blocked this response in vitro and importantly inhibited crystal-stimulated inflammation in vivo. We found in vitro and in vivo that VBY-825 acted to inhibit crystal-stimulated ASC-speck formation, and therefore was blocking the activation of the NLRP3 inflammasome. This was consistent with our earlier biochemical analyses of cultured cells (39).

We also found that MSU crystals stimulated an inflammasome-independent inflammatory response in vivo, similar to what we had earlier reported with silica (36). In this pathway, serine proteases such as protease 3, cathepsin G, and neutrophil elastase are thought to cleave extracellular pro-IL-1β to mature IL-1β (40). These proteases are synthesized as zymogens that are activated when cleaved by cathepsin C or maybe other proteases (41, 42). VBY-825 does inhibit cathepsin C at higher concentrations (Supplemental Figure 2B).

Gout is caused by the deposition of MSU crystals in tissues which then elicits a robust acute inflammatory response. In gout patients, this crystal nucleation often occurs in joints and causes severe arthritis. Administering IL-1 blocking therapy, such as Anakinra (IL-1Ra) or Canakinumab (anti-IL-1β), to such patients inhibited this arthritis (10, 11) showing that IL-1 is a key proinflammatory cytokine driving this inflammation, like it was in mouse models (7, 8). In patients, acute flares of gout can be managed by treating with non-steroidal anti-inflammatories (NSAIDs), colchicine, or corticosteroids (10, 11). However, many gout patients have contraindications against taking NSAIDs or colchicine (in 90+% or 30-40% of patients, respectively), may not tolerate these treatments, and/or may not respond to therapy (43). Our findings suggest that broadly active cathepsin inhibitors have the potential to be developed as another therapeutic to help manage these patients.

The pathogenesis of other crystal-based diseases, such as silicosis and asbestosis, is similarly thought to be caused by crystal-stimulated inflammation (3). In mouse models, the development of silicosis requires the NLRP3 inflammasome (35). Unfortunately, we have lacked therapeutics to treat and prevent the progression of these diseases. In addition, cholesterol crystals have been implicated in driving IL-1β-dependent inflammation and the development of atherosclerosis in the aorta (5, 44). Based on this a clinical trial treating patients with advanced atherosclerotic disease with Canakinumab was performed and showed benefit in reducing recurrent cardiac events and lowering C-reactive protein levels (45). Yet another disease that may be linked to the particle-stimulated inflammatory response is Alzheimer’s disease, where the particle is thought to be beta-amyloid aggregates (46, 47). Since cathepsin inhibitors can block such crystal/particle-stimulated inflammation, our findings raise the possibility that these agents might be useful in treating these diseases.

Based on the role of IL-1β in driving the above and other diseases, there has been considerable interest in developing inhibitors of the NLRP3 inflammasome. A number of small synthetic molecule or natural product inhibitors of NLRP3 inflammasomes have been identified (48). Many of these compounds are not fully specific for NLRP3 inflammasomes and can inhibit additional targets. A number of these compounds have been found to decrease, albeit partially, MSU-stimulated inflammation in mice (49) or in one case, gout arthritis measures in patients (50). Our findings, that there is a substantial inflammasome-independent pathway contributing to MSU and silica-induced inflammation, might be a reason why the inflammasome inhibitors generally achieved only a partial reduction in inflammation in vivo. Perhaps consistent with this idea, human gout patients treated with the inflammasome inhibitor Dapansutrile, did not have a reduction in serum IL-1β, despite having reduced intracellular and secreted levels of IL-1β in their PBMC (50). As noted above, cathepsin inhibitors have a theoretical advantage in that they block both the inflammasome-dependent and independent pathways and VBY-825 did achieve a strong suppression of MSU-induced inflammation and arthritis in vivo.

In summary, cathepsin inhibitors may have the potential as therapeutic agents to treat crystal and particle-based diseases. We showed proof of concept of this potential in a preclinical gout model. The cathepsin inhibitors have a different mechanism of action than existing therapeutics that could offer advantages. For example, and as noted above, they block the two major pathways of IL-1β production. Moreover, cathepsin inhibitors are relatively selective for blocking responses stimulated by sterile particles, don’t block IL-1β production from other types of inflammasomes or pro-inflammatory agents (20) and don’t block microbial endotoxin (LPS)-stimulated inflammation (Supplemental Figure 4). Consequently, these properties might not limit many of the anti-microbial defense functions of inflammatory responses, which in contrast can be interfered with by other agents, e.g., steroids or total IL-1 blockade, and thereby increase the risk of infection (51). Cathepsin inhibitors have been administered to humans and were reported to be well tolerated and safe (52, 53). In addition, these agents should be less costly than, e.g., biologics and some of the cathepsin inhibitors are orally bioavailable. Obviously, further studies will be needed to determine the potential of this class of agents as drugs.

## Methods

### Sex as a biological variable

Our study examined male and female animals, and similar effect of inhibitors were seen in both genders.

### Reagents

MSU crystals were prepared as described in the literature (7). Briefly, uric acid (5 mg/mL; Sigma-Aldrich) was dissolved in 0.1 M borate buffer on a stirrer hot plate, the solution was adjusted to pH 9 until the powder was completely dissolved and then reduced to pH 7.2. The solution was then filtered through a 0.45 µm filter (Merk-Millipore, SA1J79H5) and left in the cold room for 18 hours. The solution was again filtered through a 0.8 µm filter (Merck-Millipore SA1J95H5) and left in the cold room with steady stirring until needle-like crystals were formed (in around 2-3 days). Crystals were then collected and washed twice in acetone and once in ethanol, dried, and sterilized. Finally, the crystal size was reduced by using a focused-ultrasonicator (Covaris E220e) for 1 minute to obtain crystals <10 µm in length. They were aliquoted in sterile PBS and kept at 4°C until use. LPS was from Sigma-Aldrich. CyQUANT LDH Cytotoxicity Assay was from Invitrogen. Anti-CD11b (M1-70), anti-F4/80 (BM8), and anti-Ly6G (1A8) were purchased from BioLegend. Anti-Ly6B (7/4) was purchased from BioRad. DAPI was included for samples being analysed by Amnis FlowSight. VBY-825 was obtained as a gift from Leslie Holsinger at ViroBay. CA-074 Me (CAS No: 147859-80-1) was purchased from MedChemExpress.

### Mice

ASC-citrine transgenic, Casp1/11^-/-^ and IL-1R1^-/-^ mice were kindly provided by Dr. Douglas Golenbock (UMass Chan Medical School). Wild type (WT) C57BL/6J (B6) mice were purchased from Jackson Laboratory. All the mice used in these experiments were maintained with water and food ad libitum and 12h/12h light cycle in the animal facility (UMass Chan Medical School) and were sex- and age-matched (females 7-8-week-old or males 12-13-week-old).

### Macrophage stimulation

Murine RPMs, 1x10^5^/well, were primed with LPS (200 ng/mL) for 2 hours, followed by adding media, VBY-825 (120 µM) or CA-074 Me (5 µM) for another hour, and then media, Silica (15 µg/well), or MSU (25 µg/well) was added and cells were incubated for 4 hours. IL-1β and TNF-α in the culture supernatant was measured by ELISA kits (R&D System) according to the manuals. Cell death was measured by detecting LDH activity in the culture supernatant using CyQUANT LDH Cytotoxicity Assay (Invitrogen) according to the manufacturer’s instructions.

### Peritonitis model

Protocol for the peritonitis model was adapted from the literature (36). Briefly, mice were i.v. injected with or without VBY-825 (7.5 or 10 mg/Kg) or CA-074 Me (10 mg/Kg) 1 hour prior to i.p. injection of MSU (0.1 mg/mouse), Silica (0.1 mg/mouse), LPS (25 ng/mouse), IL-1β (1 ng/mouse) or PBS (as negative control). Four hours after particle injection, the peritoneum was lavaged with 6 ml RPMI-1640 medium with 2% FCS, 3 mM EDTA, and 10 U/ml heparin. Cells in the lavage were stained for neutrophils (Ly6B^+^, Ly6G^+^) and inflammatory macrophages (iMacrophages, Ly6B^+^, Ly6G^-^). The cell numbers were counted by using Count Bright Absolute Counting Beads (Invitrogen) and flow cytometry.

### ASC Speck analysis

RPMs (10^6^ cells) from ASC-citrine transgenic or WT mice were primed with 240 ng/ml LPS for 3 hours at 37°C in an eppendorf tube, followed by treatment with 120 µM VBY-825 or vehicle for 1 hour. Subsequently, the macrophages were stimulated with or without 100 µg of MSU or 60 µg of Silica for another 4 hours before staining them with anti-CD11b, anti-F4/80, and DAPI. The cells were analyzed by imaging flow cytometry (Amnis FlowSight) and IDEAS software (Amnis). For in vivo speck formation, ASC-citrine mice were i.v. injected with VBY-825 (7.5 mg/Kg) for 1 hour before i.p. injection with MSU (0.1 mg/mouse) for another 4 hours. The PECs were collected, stained with anti-CD11b, anti-F4/80, and DAPI, and analyzed by imaging flow cytometry.

### MSU-induced arthritis

Male mice (12–13-week-old) were depilated around the knee area using hair removal cream (Nair) 1 week before intraarticular (i.a.) injections. One hour before knee injection, mice were i.p. injected with 1 mg/mouse VBY-825 or vehicle. Knee joint inflammation was then induced by i.a. injection of MSU (150 μg/10 μL) under the patellar ligament of mice using a 10 μL Hamilton syringe with a 30G needle. Control animals received an i.a. injection of sterile saline (10 μL) under the same conditions as the MSU group. Twenty-four hours after i.a. injection, mice received i.v. Luminol (200 mg/Kg, Millipore) under isoflurane anesthesia and were immediately placed in an IVIS Spectrum CT instrument (Perkin-Elmer) and photographed. The images were analyzed using Living Image software. After IVIS imaging, mice were euthanized and the knee joints were dissected, fixed with 10 % buffered formalin for a week, and then decalcified with Decalcifier I (Leica) overnight and embedded in paraffin for histological analysis (UMass Chan Medical School Morphology Core Facility). The paraffin sections (7 µm) were stained with hematoxylin and eosin for conventional morphological evaluation using a light microscopy. Slides were assessed by 3 people and evaluated using a synovitis score modified from *V Krenn et al. 2006* (34) as following scale: 0= no change; 1= slight increase in synovial cellularity, 2= moderate increase in synovial cellularity; 3= Few perivascular neutrophils and/or mononuclear cells; 4= Many neutrophils and/or mononuclear cells; 5= dense band-line neutrophil infiltrate.

### Cathepsins inhibition assay

The enzyme activities of commercial mouse cathepsin C (mCatC, Fisher Scientific) and human cathepsin L (hCatL, R&D Systems) were quantified using AMC-based fluorogenic substrates (Z-Leu-Arg-AMC, R&D Systems, for hCatL and Gly-Arg-AMC, Cayman Chemical Company, for mCatC). For the mCatC assay, the reaction was carried out in 50 mM MES pH 5.5, 50 mM NaCl, and 5 mM DTT. Each well contains 0.2 µL of VBY-825 at varying concentrations in DMSO, followed by the addition of 10 µL 0.01 ng/µL (2x, 0.005 ng/µL final) mCatC solutions and 10 µL 200 µM Gly-Arg-AMC (2x, 100 µM final) solutions diluted from 10 mM DMSO stock with reaction buffer to initiate the reaction. Fluorescence (λex = 380 nm, λem = 460 nm) of the AMC product was monitored by Cytation 3 Multi-Mode Reader (BioTek, Winooski, VT) (54) every minute for 60 minutes at 25°C. Reaction rates were calculated using Gen 5 3.10 as RFU/sec and then normalized by the hydrolysis rates measured in the absence of VBY-825. hCatL assay follows the same procedure as the mCatC assay, except that the reaction buffer contains 50 mM MES pH 6.0, 5 mM DTT, 1 mM EDTA-Na_2_, 0.005 % (w/v) Brij-35. The final hCatL concentration is 0.02 ng/µL, and the final substrate Z-Leu-Arg-AMC concentration is 5 µM.

### Statistical analyses

In vitro data are presented as mean ± SD and In vivo data as means ± SEM. Statistical analysis was performed with an unpaired, two-tailed Student *t*-test when only two groups were compared. When more than two groups were compared, One/Two-way ANOVA and Šidák’s/ Tukey’s/ multiple comparison posttest were used to compare the means of every group to every other group. Kruskal-Wallis test followed by Dunn’s multiple comparisons test was used for non-parametric comparison. A p value <0.05 was considered statistically significant. GraphPad Prism Software was used for all the analysis.

### Study approval

All animal protocols were approved by the UMass Chan Medical School’s IACUC under protocol number 201900318.

## Data availability

Values for all data points in the graphs are reported in the Supporting Data Values file.

## Statement of author contributions

LAAO, CZ and JJL conceived the study and carried out experiments. LAAO, JJL, MB, and KLR conceived the study and analyzed the data. JZ carried out experiments. LAAO, JJL and KLR wrote the manuscript. All authors were involved in reviewing the manuscript and had final approval of the submitted version.

## Acknowledgments

This work was supported by NIH grant AI173076. We also thank assistance from the UMASS Flow Core, Morphology Core, and Animal Medicine in this study.

