## Supplemental data for "A broad cathepsin inhibitor blocks crystal-stimulated inflammasome-dependent and -independent inflammation, and gout arthritis"

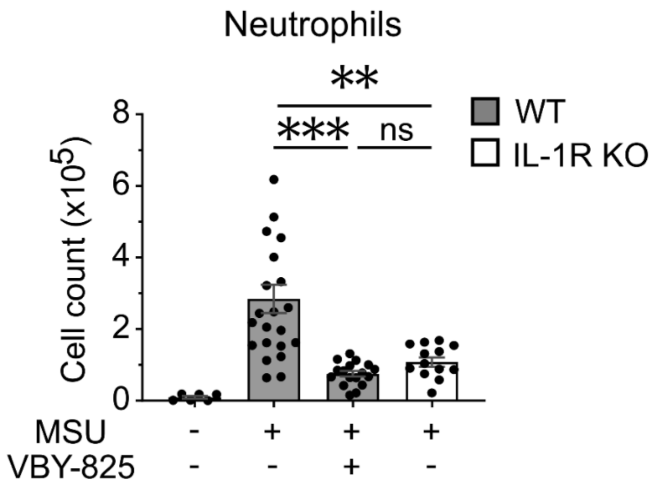


**Supplemental Figure 1. IL-1R deficiency impairs the inflammation induced by MSU crystals**

WT and IL-1R knockout (IL-1R KO) female mice were i.v. injected with or without VBY-825 (7.5 mg/Kg) for 1 hour, followed by i.p. injection of MSU crystals (0.1 mg/mouse) for another 4 hours. The mouse peritoneum was lavaged and the infiltrating neutrophils were counted by flow cytometry. The data were combined from 3 experiments and represented as means ± SEM. **p<0.01; ***p<0.001.

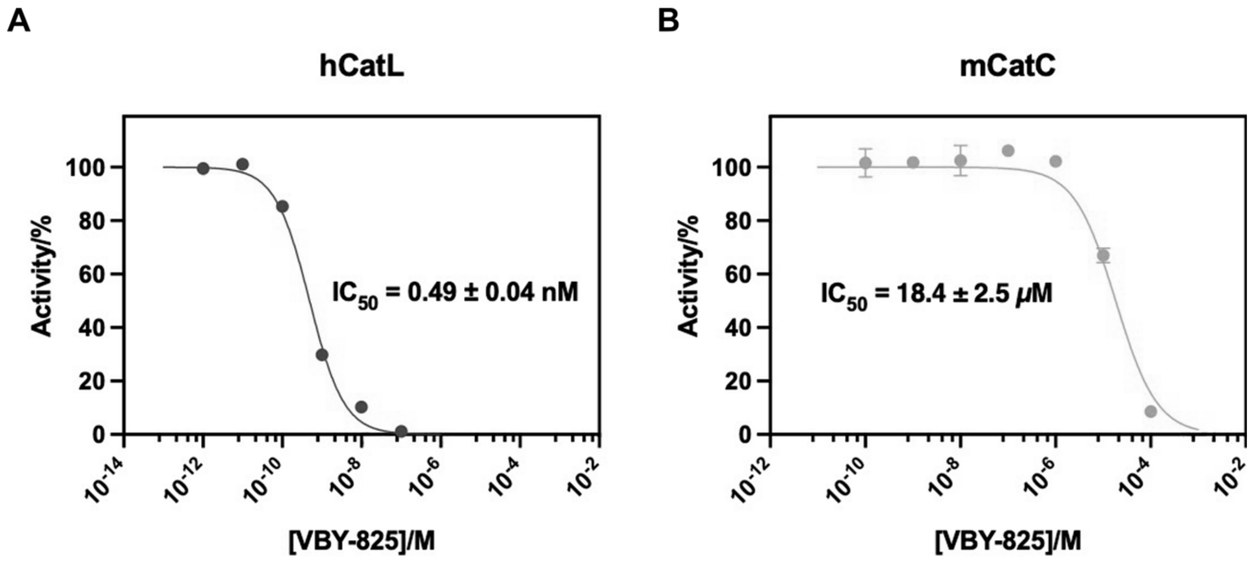


**Supplemental Figure 2. VBY-825 suppresses the enzyme activity of Cathepsin L and Cathepsin C**

Dose-response curves to determine the $IC_{50}$ of compound VBY-825 against recombinant human cathepsin L (**A**) and mouse cathepsin C (**B**). $IC_{50}$ values were calculated from the percentage of enzyme activity with data points and bars representing means ± SD (*n* = 2).

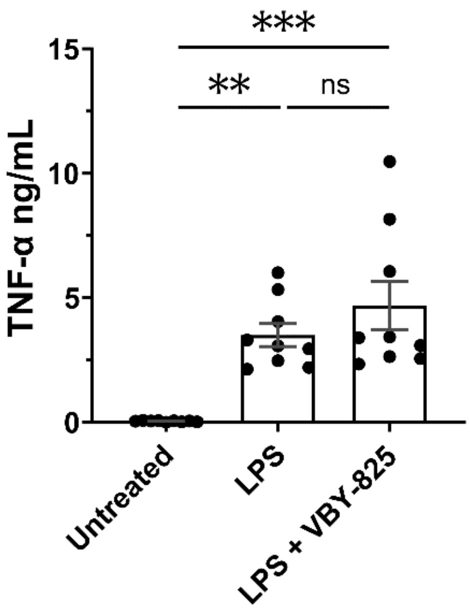


**Supplemental Figure 3. VBY-825 does not suppress TNF-α production induced by LPS in macrophages**

RPMs, $1x10^5$/well, were incubated with LPS (200 ng/mL) for 6 hours with or without pre-treatment with VBY-825 (120 µM) for 1 hour. TNF-α in the culture supernatant was measured by ELISA. The data are combined results of 3 experiments and represented as means ± SD. **p<0.01, ***p<0.001.

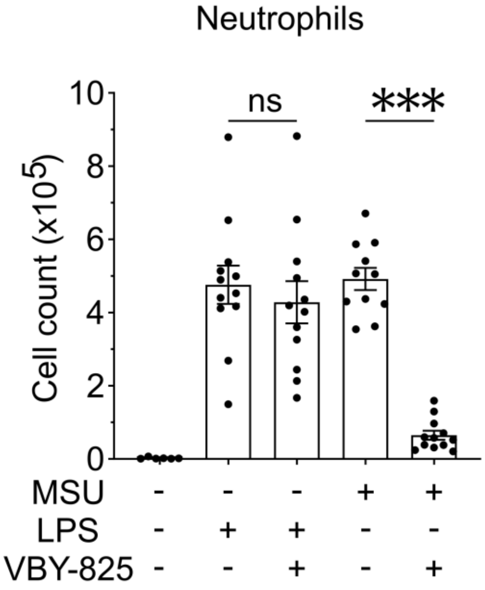


**Supplemental Figure 4. VBY-825 does not suppress peritonitis induced by LPS**

WT female mice were i.v. injected with or without VBY-825 (7.5 mg/Kg) for 1 hour, followed by i.p. injection with or without MSU crystals (0.1 mg/mouse) or LPS (25 ng/mouse) for another 4 hours. The neutrophils in the peritoneum were lavaged and counted by flow cytometry. The data are combined results of 3 experiments and represented as means ± SEM. ***$p < 0.001$.
